# Microsaccades strongly modulate but do not cause the N2pc EEG marker of spatial attention

**DOI:** 10.1101/2024.10.28.620656

**Authors:** Baiwei Liu, Siyang Kong, Freek van Ede

## Abstract

The N2pc is a popular human-neuroscience marker of covert and internal spatial attention that occurs 200-300 ms after being prompted to shift attention – a time window also characterised by the spatial biasing of microsaccades. Here we show how co-occurring microsaccades profoundly modulate N2pc amplitude during the top-down deployment of spatial attention in both perception and working memory. At the same time, we show that a significant – even if severely weakened – N2pc can still be established in the absence of co-occurring microsaccades. Thus, while microsaccade presence and direction strongly modulate N2pc amplitude, microsaccades are not strictly a prerequisite for the N2pc to be observed.

## Introduction

Attention is the foundational process by which we selectively process and prioritize information that is relevant to us ^1,2^, a central topic within the study of mind and brain. Attention can be deployed covertly to locations outside of current fixation, as well as internally to representations held in working memory ^3,4^. A widely used marker of covert and internal attention shifts in human neuroscience is the N2pc – a lateralized posterior EEG potential that emerges around 200 ms after being prompted to shift attention. Since the discovery of the N2pc around 30 years ago ^5^, this marker has provided invaluable insights into the principles, mechanisms, and timings of covert and internal attention shifts ^5–15^.

A vital assumption in laboratory studies of the N2pc is that participants remain their current fixation while shifting attention. Only then can the spatial modulation of brain activity, like the N2pc, be attributed to a top-down cognitive modulation associated with attention (as opposed to consequences of eye-movements or the retinal displacements they trigger). The validity of this vital assumption, however, is not guaranteed. Even when instructed to keep fixation, humans nevertheless produce small – often overlooked – involuntary eye movements known as microsaccades ^16–18^. Critically, the direction of microsaccades has also been shown to be modulated by the top-down deployment of covert ^19–21^ and internal ^21–26^ spatial attention. Moreover, spatial modulations in microsaccades during covert and internal attention shifts occur at time windows that overlap with the N2pc, also emerging around 200 ms after being prompted to shift attention. This raises the critical question to what extent the N2pc is contingent on – or even caused by – co-occurring spatial biases in microsaccades.

Despite the extensive use of the N2pc for studying top-down attention, the potential contribution of microsaccades to the N2pc remains elusive (unlike for complementary neural attention markers ^22,27,28^ or spatial ERP modulations associated with bottom-up cue processing ^29^). Moreover, prior studies on the link between microsaccades and attention almost exclusively considered covert shifts of spatial attention in the context of perception. Here, microsaccades may also influence neural activity through retinal displacement of the attended sensory inputs. In contrast, when shifting attention to visual contents held in visual working memory, the potential contribution of retinal displacement is sidestepped because the attended objects are in memory. To address these outstanding points, we investigated the contribution of microsaccades to the N2pc and studied this during the top-down deployment of spatial attention in both perception and in working memory.

## Results

Healthy human volunteers participated in two complementary attention tasks that either required covert selection of one of two visual objects displayed to the left or right on the screen (**Fig. 1a**), or internal selection of either of these visual objects from working memory (**Fig. 1h**). In both cases, attentional selection was prompted via a central, non-spatial, color cue. Participants were required to compare the cued (color-matching) visual object to an ensuing centrally presented black probe stimulus, judging whether it was rotated clockwise or counter-clockwise. Note how in this set-up the use of the cue was imperative: without using the cue participants would not know to which visual object the probe should be compared. Participants were well able to perform both the perceptual (reaction time: 908 ± 60 ms (M ± SE); accuracy: 90 ± 1 %) and working-memory (reaction time: 922 ± 61 ms; accuracy: 84 ± 1 %) versions of this selective attention task.

**Figure 1.**
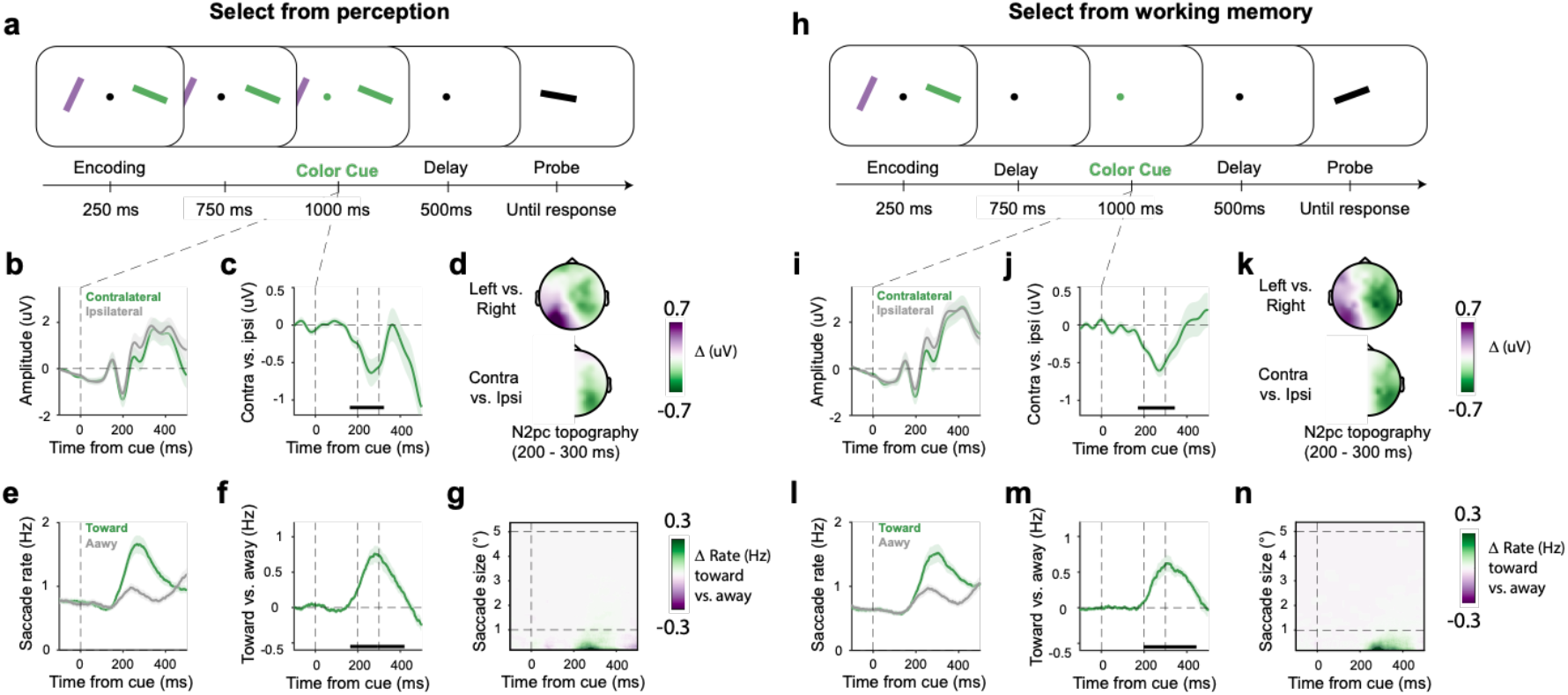
Attentional selection evokes an N2pc and biases microsaccade directions at overlapping time windows in both perception and working memory. Participants performed two tasks requiring covert selection of visual objects from perception (**a-g**) or working memory (**h-i**) following a central color cue. **a, h**) Task schematics. In both versions of the task, an imperative central color cue indicated which visual objects had to be selected to compare its orientation to the upcoming probe stimulus. **b, j**) Grand-averaged ERP waveforms in posterior electrodes PO7/8 contralateral and ipsilateral to the side of the cued object. **c, j**) The N2pc waveform, reflecting the difference between contralateral and ipsilateral ERPs. **d, k**) Topographies showing activity following cues prompting selection of the left vs. right object (top), and collapsed to show contra vs. ipsilateral (bottom). **e, l**) Time courses of saccade rates (number of saccades per second) for saccades toward and away from the side associated with the cued object. **f, m**) The spatial saccade bias, plotted as the difference between the rate of toward and away saccades. **g, n**) Difference in saccade rates (toward minus away) as a function of saccade size. For reference, dashed horizontal lines indicate a common threshold for microsaccades (1° visual angle), as well as the centre location of the selected object at 5° visual angle. All time courses show mean values, with shading indicating ±1 SEM calculated across participants (n = 23). Black horizontal lines in the time course plots indicate the significant temporal cluster (cluster-based permutation ^30^).

### Attentional selection evokes an N2pc and a spatial microsaccade bias at overlapping time windows in both perception and working memory

Following the cue to covertly select the cued visual object from perception, we were able to establish the classic N2pc marker of spatial attention shifts: event-related potentials (ERPs) were more negative contra-compared to ipsilateral to the side of the cued object (**Fig. 1b**), as quantified in predefined N2pc electrodes (PO7/PO8). **Figure 1c** zooms in on the spatial N2pc, displayed as a difference wave. Cluster-based permutation analyses confirmed a clear spatial modulation (cluster P < 0.001), with a cluster ranging from 162 to 323 ms after cue onset. Consistent with a modulation of visual-spatial attention, this difference wave had a predominantly posterior topography (**Fig. 1d**).

In a similar time window where we found N2pc, we also observed a robust bias in the direction of eye-movements according to the side of the to-be-attended stimulus (**Fig. 1e-g**). Following the cue, we found a higher rate of saccades toward vs. away from the cued object (**Fig. 1e**). The spatial saccade bias, again quantified as a difference wave (**Fig. 1f**), was also highly significant (cluster P < 0.001) with a cluster ranging from 164 to 421 ms after cue onset. By visualising this spatial bias as a function of saccade size (**Fig. 1g**), we confirmed that this bias was driven almost exclusively by small saccades (<1°) in the classical microsaccade range (objects in the display were centred at 5°), consistent with prior studies during covert and internal shifts of attention ^19–22,26^. This minute nature of the saccades driving this bias is one reason why such biases are easily overlooked in studies on the N2pc.

Highly similar results were found in the complementary task where the cue came *after* the visual objects were no longer visible – therefore prompting attentional selection within the spatial layout of working memory (**Fig. 1h-n**). Following the retro-cue, we again found a clear N2pc (cluster P < 0.001) (consistent with ^6,11^) whose significant cluster ranged from 170 to 343 ms (**Fig. 1j**), and that again had a posterior topography (**Fig. 1k**). We also again found a clear spatial saccade bias (cluster P < 0.001) whose significant cluster ranged from 198 to 445 ms (**Fig. 1m**), and that was again driven by eye-movements in the microsaccade range (**Fig. 1n**).

In both tasks we thus observed not only clear N2pc EEG components associated with top-down shifts of spatial attention, but also robust spatial biases in microsaccades that co-occur in time. Having established this vital starting point, we now turn to our core question: how are these two markers of spatial attention related?

### Microsaccades strongly modulate but do not cause the N2pc EEG marker of top-down covert and internal shifts of spatial attention

To investigate the dependence of the N2pc on the co-occurring spatial biasing of microsaccades, we adopted an approach also used and outlined in ^22^. We leveraged the fact that, unlike continuous EEG signals, microsaccades are discrete events that can be classified at the single trial level. To sort trials into relevant microsaccade classes, we defined the relevant attention window as the time window from 150 to 400 ms after cue onset (we chose this window to encompass microsaccades that slightly preceded or followed the N2pc). Having set this window, we defined three classes of trials: trials with (1) a microsaccade in the direction of the cued visual object (“toward-microsaccade trial”), (2) a microsaccade in the opposite direction (“away-microsaccade trial”), or (3) no discernible microsaccade anywhere in this key time window of interest (“no-microsaccade trial”).

This analysis confirmed a dominance of toward vs. away microsaccades (consisted with the spatial bias reported and quantified above), but also revealed a substantial proportion of trials in which no discernable microsaccade was detected in the window of interest (**Fig. 2e**). During perceptual selection, 37.5 ± 2% (M ± SE) of trials were classified as toward-microsaccade trials, 23.9 ± 3% as away-microsaccade trials, and 38.6±3% as no-microsaccade trials. Similarly, during working-memory selection, 34.9 ± 2% of trials were classified as toward-microsaccade trials, 24.5 ± 1% as away-microsaccade trials, and 40.7 ± 3% as no-microsaccade trials. This enabled us to separately examine the N2pc as a function of microsaccade class, with ample trials in each class.

**Figure 2.**
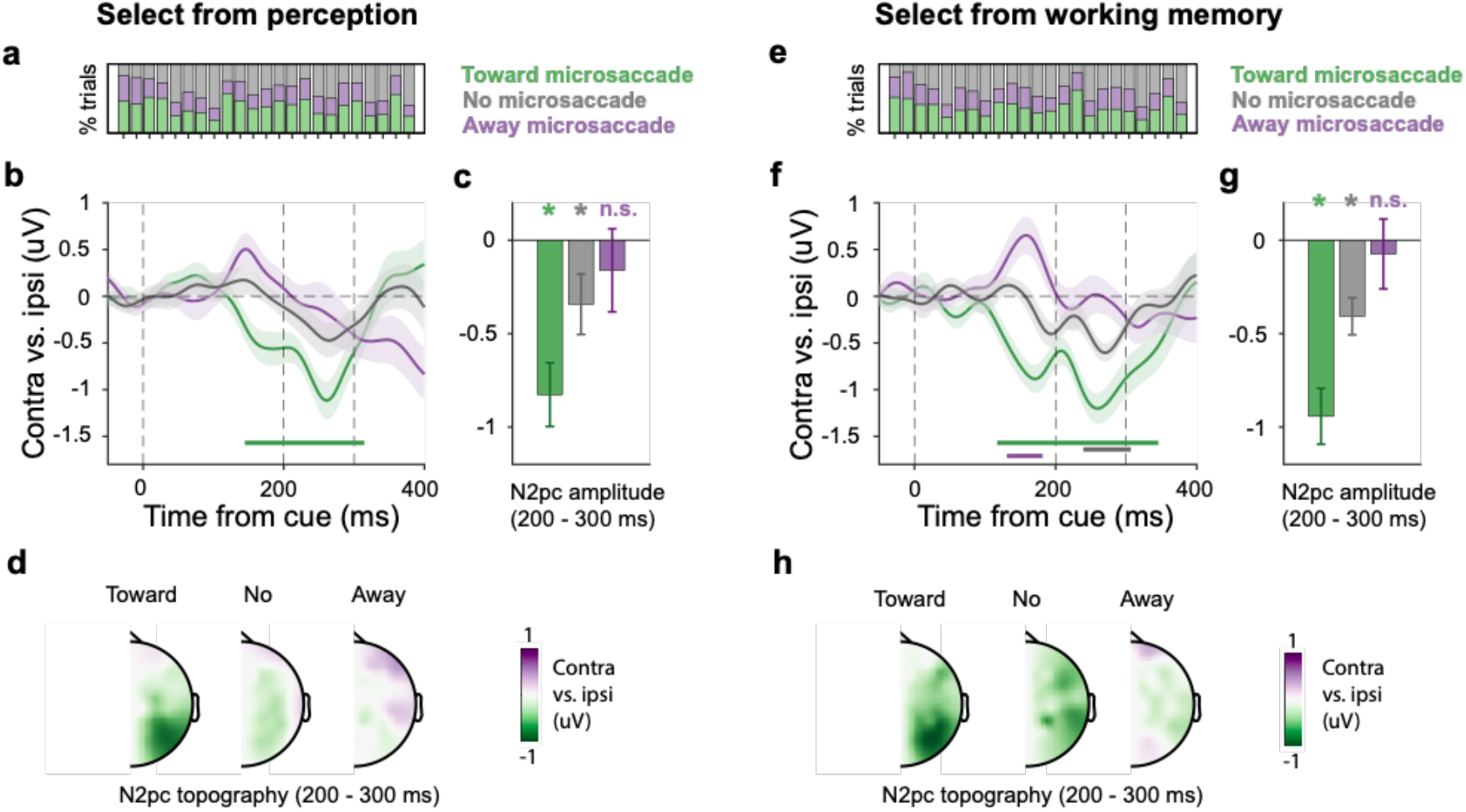
Microsaccades strongly modulate but do not cause the N2pc EEG marker of top-down covert and internal shifts of spatial attention. Trials were classified into “toward-microsaccade”, “away-microsaccade”, and “no microsaccade” classes based on the presence/absence and direction of microsaccades in the relevant predefined 150-400 ms time window. **a, e**) Participant-specific proportions of trial classes during selection from perception (panel **a**) and working memory (panel **e**). **b, f**) N2pc waveforms (contra vs. ipsilateral ERPs in PO7/8) for the three microsaccade-based trial classes. Shading indicates ±1 SEM calculated across participants (n = 23). Colored horizontal lines in the time course plots indicate significant temporal clusters (cluster-based permutation). **c, g**) N2pc amplitudes in the three microsaccade trial classes extracted over the predefined 200–300 ms N2pc time window. The bar graphs show mean values, with error bars indicating ±1 SEM calculated across participants (n = 23). The stars above the bars indicate that N2pc amplitude for that specific trial class is significantly different from zero (P < 0.05). **d, h**) N2pc topographies across trial classes plotted as contralateral vs. ipsilateral activity.

When selecting from perception, trials with a toward microsaccade showed by far the clearest N2pc (**Fig. 2b** green line; cluster P < 0.001), with a posterior topography (**Fig. 2d**). In contrast, no N2pc cluster was found in neither the away-microsaccade nor in the no-microsaccade trial-classes (**Fig. 2b**, purple and grey lines).

To increase sensitivity, we also zoomed in the a-priori defined N2pc time window from 200 to 300 ms after cue onset (**Fig. 2c**) – a window consistent with ample prior studies reporting an N2pc in this time range ^5,7–10,12–15,31^ as well as with the data from our tasks (see **Fig. 1c,j**). A one-way ANOVA revealed a strong effect of microsaccade type (*F*(2, 44) = 6.92, *P* = 0.002, partial *η*^*2*^ = 0.239). When considering individual conditions, we confirmed a highly robust N2pc in toward-microsaccade trials (t(22) = -4.82, P < 0.001, d = -1.01) and now also recovered a weak but significant N2pc in the no-microsaccade trials (t(22) = -2.12, P = 0.046, d = -0.44). However, even after averaging across the relevant N2pc time window, we still could not establish a significant N2pc in away-microsaccade trials (t(22) = -0.724, P = 0.447, d = -0.15). Direct comparisons further confirmed that the N2pc amplitude in toward-microsaccade trials was significantly larger than in both no-microsaccade (t(22) = -3.12, P_Bonferroni_ = 0.015, d = -0.65) and away-microsaccade trials (t(22) = -3.07, P_Bonferroni_ = 0.017, d = - 0.64). No significant difference in N2pc amplitude was observed between no-microsaccade and away-microsaccade trials (t(22) = -1.02, P_Bonferroni_ = 0.96, d = -0.21).

Note how these differences in N2pc across the three microsaccade classes are unlikely accounted for by mere differences in the number of available trials. For example, while the N2pc more than halved in size when moving from toward-to no-microsaccade trials, we had in fact more (not less) trials in the latter condition.

A strikingly similar pattern of results was found when the cue prompted selection from working memory (**Fig. 2e-h**). We again found the clearest N2pc in toward-microsaccade trials (cluster P < 0.001). This time we also found a significant N2pc-cluster in the no-microsaccade trials (cluster P = 0.005). However, in the away-microsaccade trials, we only observed an earlier cluster (cluster P = 0.03) associated with a reversed ERP lateralisation. When again zooming in on the N2pc time window – our key focus here – (**Fig. 2g**), we again found a strong main effect of microsaccade trial-class (*F*(2, 44) = 7.86, *P* = 0.001, partial *η*^*2*^ = 0.263). When considering individual conditions, we again observed a robust N2pc in toward-microsaccade trials (t(22) = -6.31, P < 0.001, d = -1.32) and a smaller but still robust N2pc in the no-microsaccade trials (t(22) = -4.11, P < 0.001, d = -0.86). However, just like we reported during perceptual selection, also during mnemonic selection we could not establish a significant N2pc in away-microsaccade trials (t(22) = -0.39, P = 0.7, d = -0.08). Direct comparisons again showed that N2pc amplitude in toward-microsaccade trials was significantly larger than in both no-microsaccade (t(22) = -2.92, P_Bonferroni_ = 0.024, d = -0.61) and away-microsaccade trials (t(22) = -3.22, P_Bonferroni_ = 0.012, d = -0.67). Again, no significant difference was observed between no-microsaccade and away-microsaccade trials (t(22) = -1.66, P_Bonferroni_ = 0.33, d = -0.35).

To ensure that the absent N2pc in the away-microsaccade trials was not due to a general failure to shift attention in these trials, we finally examined two additional aspects of our data: behavioral performance and the lateralization of 8-12 Hz alpha-band activity which is another canonical neural marker of spatial attention shifts ^22,32–34^. First, participants responded accurately (all conditions >80% accuracy) and quickly (all conditions <1000 ms) across toward, away, and no-microsaccade trials in both the perceptual-selection and working-memory selection tasks (**Supplementary Fig. 1**). If participants simply did not shift attention in the away-microsaccade trials, we should have seen strongly impoverished performance (the imperative cueing procedure we used that necessitated using the cue to perform the task). Second, we observed clearly preserved alpha-band lateralization reflecting spatial attention shifts in all trial classes (**Supplementary Fig. 2**). This provides direct evidence that spatial attention was also deployed in away microsaccade trials, even if we could not establish an N2pc in these trials. These data further show how the lack of an N2pc in these trials is not simply owing to a lack of sensitivity, or poor EEG signal in these trials.

## Discussion

As our data make clear, voluntary shifts of spatial attention in perception and working memory are not only associated with the N2pc (as also in ^5–14^) but also with spatial biases in microsaccades (as also in ^19–22,26^) that co-occur in time. We now further show how co-occurring microsaccades profoundly modulate N2pc amplitude during the top-down deployment of spatial attention – even if a severely weakened N2pc can still be observed in the absence of co-occurring microsaccades.

As good news for the field, we show that microsaccades do not *cause* the N2pc: the N2pc can still be observed in the absence of co-occurring microsaccades, albeit weak. This resonates with recent studies on complementary neural markers of attention (cf. ^22,28,29^; but see also ^27^). At the same time, we unveil that co-occurring microsaccades matter a lot for the N2pc: the N2pc more than doubles in size when microsaccades co-occur in the attended direction, while the N2pc vanishes when microsaccade co-occur in the opposite direction. This makes clear how microsaccades cannot be shoved under the carpet when studying this popular human-neuroscience marker of attention. Admittedly, we studied the N2pc and microsaccades in relatively simple tasks that (1) enabled us to match the perceptual and working-memory selection tasks and (2) we knew would yield both markers. Prompted by our findings, it will now be vital for the field to adopt similar analyses in complementary tasks in which the N2pc is frequently reported.

One interpretation for the reported reduction and even lack of the N2pc in trials with no or away microsaccades is that attention may simply not have been deployed to the same extent in these trials. Additional features of our dataset counter this interpretation. First, because of the way we designed the task, if participants did not use the cue they would have had no way of knowing which object was tested. This would have resulted in a large performance drop in these trials. We found no evidence for this. Second, a complementary marker of spatial attention – the lateralisation of posterior alpha-band activity ^22,32–34^ – was observed in all three microsaccade trial classes. Critically, this marker was robust even in the away-microsaccade trials where we no longer found an N2pc. These data suggest that the reported N2pc modulations are not due to differences in the degree of attentional deployment per se, but to the presence and direction of co-occurring microsaccades. Further note how the microsaccade-class-specific differences in N2pc amplitude are also unlikely accounted for by differences in trial numbers. For example, while the N2pc more than halved in size when moving from toward-to no-microsaccade trials, we had in fact more (not less) trials in the latter condition.

Our work complements prior studies linking microsaccades to neural activity ^35–40^ and to neural modulations by spatial attention ^22,27–29^. We advance the latter literature in two critical ways. First, we uniquely targeted the N2pc, a canonical ERP marker of voluntary spatial attention from the human-neuroscience literature. While at least one prior study ^29^ also considered spatial ERP modulations to attention cues, ERP lateralisation and microsaccade biases in that study reflected bottom-up processing of the spatial cue, not top-down deployment of attention. Because we used a central color cue, any spatial modulation in our task must reflect top-down shifts of attention – the type of attention classically studied with the N2pc. Second, we not only considered attention in the context of perception (as in most prior studies) but also working memory. This not only offered an internal replication, but also brings an important advance. This shows how the reported N2pc modulations by microsaccades hold regardless whether microsaccades also bring attended visual targets closer to the fovea (in perception) or not (in working memory).

By measuring EEG, our data only allow us to conclude about N2pc presence and N2pc magnitude from the view of extracranial neural measurements. The fact that we did not observe an N2pc in away microsaccade trials at the scalp level, does not necessarily imply that no N2pc occurred anywhere inside the brain, as the N2pc may have been “masked” by additional signals imposed by the microsaccade. This possibility of ‘signal mixing’ is hard to address based on the current data and likely requires intracranial measurements ^41^ or MEG ^42^, as well as additional types of analyses. This was beyond the scope of our current study, which was to delineate how much microsaccades contribute to the N2pc – both in perception and working memory – when following procedures that are custom in N2pc research.

Finally, we appreciate how some may view microsaccades as a threat to the study of covert and internal attention by virtue of their confounding contribution to neural markers like the N2pc. However, microsaccades can also be seen as an opportunity ^26,43^. Like the N2pc, microsaccade biases also enable tracking of spatial attention shifts; they too have high temporal resolution; and they too track spatial attention shifts in both perception ^19,20^ and working memory ^22–25^. In future studies it will therefore be interesting not only to systematically compare the inter-relation of both markers, but also their respective usefulness for studying covert and internal shifts of attention across different experimental tasks and settings.

## Methods

### Ethics

Experimental procedures were reviewed and approved by the local Ethics Committee at the Vrije Universiteit Amsterdam. Participants provided written informed consent prior to the experiment and were compensated €10 per hour (or the equivalent in credits) for their time.

### Participants

Twenty-three healthy human volunteers participated in the study (age range: 19-32; 2 male and 21 female; 22 right-handed; 15 corrected-to-normal vision). The sample size of 23 was set a-priori based on a prior study from the lab that addressed a similar research question ^22^ for a complementary neural signature. To reach the desired sample size, the first participant had to be replaced due to an error in the experiment code.

### Task design and procedure

Participants performed two complementary spatial-attention tasks that either required covert selection of one of two visual objects displayed to the left or right on the screen (select from perception) or internally select either of these visual objects that were no longer visible at the time of the attention cue (select from working memory; **Fig. 1h**). We designed our tasks in such a way that they were as similar as possible, and only differed whether task performance relied on visual object selection from perception or from working memory.

Trials started with the presentation of two visual objects (bars with distinct colors and randomly drawn orientations) centred at 5 degrees to the left and right of a central fixation dot. Bars were 2 by 0.4 degrees visual angle, and the fixation dot had a radius of 0.07 degree visual angle. In the perceptual-selection task (**Fig. 1a**), the stimuli stayed on the screen for 2000 ms. While the stimuli were still visible (1000 ms after stimuli onset), the central fixation dot changed color for 1000 ms, acting as a 100% valid attention cue. This cue instructed participants to select the color-matching target object for the ensuing test. After the offset of the stimuli, a blank display was presented for 500 ms, before the onset of the test display. This ensured that participants needed to select and remember the cued bar ahead of time. In the final test display, a black bar appeared at the center of the screen, rotated between 10 to 20 degrees clockwise or counter-clockwise relative to the orientation of the cued bar. Participants were required to indicate whether the cued object (meanwhile in memory) should be rotated clockwise or counter-clockwise to match the black bar in the test display. Note how cues were imperative in this task as the test display itself did not contain any information as to which bar was being tested. After responding, participants received immediate feedback indicated by a number (“0” for incorrect and “1” for correct) displayed for 250 ms just above the fixation point. Inter-trial intervals were randomly drawn from 500 to 1000 ms.

In the working-memory-selection version of this attention task (**Fig. 1h**), the procedure was identical except that the stimuli only briefly appeared at the start of each trial for 250 ms and thus were no longer visible when the cue was presented. Instead, the cue now instructed which colored object to select from working memory. The onset timings of the stimuli, the cue, and the test display were identical between the two tasks. Note also how both tasks eventually relied on a memory-guided report. The only and key difference was whether the cue prompted selection of a visual object that was currently visible (select from perception) or not (select from working memory).

In each trial, the bars were randomly assigned two distinct colors selected from a set of four: blue (RGB: 21, 165, 234), orange (RGB: 234, 74, 21), green (RGB: 133, 194, 18), and purple (RGB: 197, 21, 234). The orientations of the bars was randomly drawn from 0° to 180° with a minimum difference of 20° between them. Each bar was equally likely to be cued for report. Across trials we made sure to equally often cue the left or the right object. During the test display, the black bar (RGB: 64, 64, 64) was oriented either clockwise or counter-clockwise from the cued target object, with a random change in orientation ranging from 10 to 20 degrees.

The experiment consisted of 3 consecutive sessions, each containing 10 blocks of 40 trials. Within each session, 5 blocks contained the perceptual-selection version of our task, and 5 blocks the working-memory-selection version, in random order. The task-version was explicitly stated prior to each block. Each participant completed a total of 1200 trials, evenly divided between the perceptual (600 trials) and working-memory (600 trials) attention tasks. Prior to the formal testing, participants practiced for approximately 10 minutes to familiarize themselves with both task types.

The experiment was programmed in Python (version 3.6.13) with psychopy (version 2021.2.2). During the experiment, participants sat in front of a monitor (with a 100-Hz refresh rate) at a viewing distance of ∼70 cm with their head resting on a chin rest.

### Eye-tracking acquisition and pre-processing

We tracked the right eye for all participants with an EyeLink 1000 system (SR Research) at a sampling rate of 1000 Hz. The eye-tracker camera was positioned approximately 5 cm in front of the monitor, 65 cm away from the participant. Gaze position was continuously monitored along both the horizontal and vertical axes. Prior to recording, we used the built-in calibration and validation protocols of the EyeLink software. After recording, the eye-tracking data were converted from the original .edf format to .asc format and then analyzed in MATLAB using the FieldTrip analysis toolbox with custom scripts. Before turning to saccade-detection, we identified blinks by detecting 0 clusters in the gaze data, and then we set all data from 100 ms before to 100 ms after the detected 0 clusters to Not-a-Number (NaN) to remove any residual blink artifacts. Finally, the data were epoched from -1000 ms to +1500 ms relative to the onset of the attention cue.

### Microsaccade detection

To identify microsaccades, we employed a velocity-based detection method, which we extensively validated in our previous article addressing a similar question ^22^. Because the objects were always horizontally arranged (one left, one right), our analysis focused on the horizontal channel of the gaze data, which is also the same as in our previously validated approach.

We first calculated the gaze velocity by taking the Euclidean distance between temporally successive gaze-position values in the horizontal axis. To increase the SNR, we smoothed velocity by applying a Gaussian-weighted moving average filter with a 7-ms sliding window using MATLAB’s built-in “smoothdata” function. We identified saccades when the velocity exceeded a trial-based threshold of 3 times the median velocity, marking the first sample that crossed the threshold as the onset of the saccade. We note how the default setting in our custom saccade-detection function is a threshold of 5 times the median gaze velocity. However, for the purpose of the current question, we deliberately used a lower threshold. Doing so, we increase confidence that our “no-microsaccade” condition really did not contain any microsaccade (at the expense of the occasional misclassification of noise as a “toward-microsaccade” or “away-microsaccade” microsaccade). We adopted the exact same logic and threshold in ^22^. To prevent counting the same saccade multiple times, we imposed a minimum delay of 100 ms between successive saccades. The magnitude and direction of each saccade were determined by comparing pre-saccade gaze positions (−50 to 0 ms before saccade onset) with post-saccade gaze positions (50 to 100 ms after saccade onset). We only considered gaze shifts with an estimated magnitude of at least 0.05 visual degrees (3 arcmin), because it is hard to ascertain the direction of the smaller gaze shift. Trials in which we observed gaze shifts smaller than 0.05 degrees in the analysis window of interest were removed from further analysis.

We calculated gaze shift rates (measured in Hz) using a sliding time window of 50 ms, advanced in steps of 1 ms. To quantify the spatial biasing of saccades, we classified saccades as ‘toward’ or ‘away’ from the cued visual object (or its memorized location) and compared the rates of toward and away saccades. In addition, to verify the microsaccadic-nature of the reported spatial saccade biases, we chose not to set an arbitrary threshold on saccade sizes, but rather to visualize the spatial saccade bias as a function of saccade size (as in ^22^). For this, we decomposed shift rates into a time-size representation (**Fig. 1 g, n**). For saccade-size sorting, we employed successive saccade-size bins of 0.2 visual degrees, with increments of 0.04 visual degrees.

### Microsaccade-based trial sorting

A key step in our analysis was to separate the trials based on the presence/absence and direction of microsaccades occurring in the ‘attention window of interest’. WE set this window to 150-400 ms after cue onset. This ensured to capture the N2pc time window (from 200-300 ms post cue onset), while also including potential saccades that preceded or followed in time. If no gaze shift was detected in this key window, the trial was labeled as a “no-microsaccade” trial. If a gaze shift *was* detected, we further categorized the trial by the direction of the first detected shift in this window, classifying the trial as either a “toward-microsaccade” or “away-microsaccade” trial, in reference to the left/right location of the cued visual object.

Before trial classification, we excluded any trials where the eye-tracking data contained NaN values (typically due to blinks or temporary loss of eye signal) anywhere within attention window of interest (150-400 ms after cue onset), or trials in which the detected gaze shifts were smaller than 0.05°.

### EEG acquisition and pre-processing

We used the BioSemi ActiveTwo System (biosemi.com) with a conventional 10–10 System 64-electrode setup. Two electrodes on the left and right mastoid were used for offline re-referencing of the data. For the purpose of data cleaning, we also measured EOG, with two electrodes placed horizontally near the left and right eyes and two electrodes above and below the left eye. EOG measurements did not serve to track eye-movements (for this we used the dedicated Eyelink eye-tracker), but exclusively served as useful signals to identify ICA components for removal.

We used MATLAB (2022a) to analysis EEG with the FieldTrip toolbox ^44^ and custom codes. We first epoched the data from 1000 msec before to 1500 msec after cue onset, re-referenced the data to an average of the mastoid electrodes. After that, we conducted an Independent Component Analysis (ICA) and correlated the ICA components with recorded EOG to identify components for rejection. Finally, we removed the trials with exceptionally high variance using the function ft_rejectvisual in Fieldtrip. All the data cleaning was performed without knowledge of the experimental conditions to which individual trials belonged. All analyses were performed on data at a 1024 Hz sampling rate.

### N2pc analysis

The epoched EEG data was first baseline corrected by subtracting the average potential in the 250-msec window preceding cue onset, and then averaged across trials for each condition. To extract the N2pc, we focused on activity in predefined posterior visual electrodes PO7/PO8 (as also in ^8,15,31,45^) and calculated the difference between contralateral and ipsilateral waveforms, relative to the location (or memorized location) of the cued object. In addition to visualising and evaluating the full time courses of this ERP lateralization, we also extracted the average N2pc amplitude by averaging over the predefined window of 200-300 ms after cue onset, based on ample prior reports of the N2pc in a similar time range ^5,7–10,12–15,31^. Trial-averaged ERPs were smoothed using as Gaussian kernel with a standard deviation of 15 samples (∼15 ms at our sampling rate of 1024 Hz).

### Time-frequency analysis

Though our focus was on the N2pc (an ERP component characterized in the time domain), for completeness we also considered the lateralization of 8-12 Hz alpha-band activity. To this end, we first applied a time-frequency decomposition using a short-time Fourier transform with Hanning-tapered data. A 300-ms sliding time window was used to estimate spectral power to estimate spectral power within the 3 to 40 Hz range in the step of 1 Hz, progressing in steps of 20 ms. To quantify lateralization, we used the same predefined posterior visual electrodes (PO7/PO8) that we used to extract the N2pc. Lateralization was again calculated by comparing activity contralateral versus ipsilateral to the location of the cued visual object. For spectral power, this contrast was expressed as a normalized difference: [(contra-ipsi)/(contra+ipsi)] × 100.

### Analysis of behavioural performance

We analyses both task accuracy (percentage correct responses) and response times (counted from probe onset). Response times were cleaned using a two-step trimming procedure. First, trials with response times exceeding 3000 ms were excluded. Second, data were further trimmed using a cutoff of 2.5 standard deviations from the participant’s mean. All behavioral performance analyses were conducted on the trimmed dataset.

### Statistical analysis

For statistical evaluation of the time-series data, we used a cluster-based permutation approach ^30^, that evaluates the reliability of neural and gaze patterns across neighboring time points while controlling for (i.e., effectively bypassing) multiple comparisons. First, a permutation distribution of the largest clusters was created by randomly permuting the trial-averaged condition-specific data at the participant level. We then calculated the p-value for each observed cluster by using the proportion of permutations where the largest cluster exceeded the size of the observed cluster. We performed 10,000 permutations and identified clusters using Fieldtrip’s default settings (grouping adjacent same-signed data points significant in a mass univariate t-test with a two-sided alpha level of 0.05, and defining cluster size as the sum of all t values within the cluster).

To statistically confirm that N2pc amplitude was modulated by microsaccade trialclass (i.e., toward, away, no microsaccade trials), we additionally employed a one-way ANOVA on N2pc amplitude averaged across the predefined N2pc time window. We supplemented this with post-hoc condition comparisons, and also compared the N2pc within each trial-type to zero, to test for evidence on the absolute presence/absence of the N2pc. We performed all analyses separately for the two versions of our task in which the cue prompted selection either from perception or from working memory. A direct comparison between tasks was beyond the scope of our central aims. We included both tasks only to seek generalization across these two complementary task settings that have each been shown to yield both N2pc and microsaccade modulations ^5,6,11,19,20,22,26^.

## Acknowledgements

This work was supported by an ERC Starting Grant from the European Research Council (MEMTICIPATION, 850636) and an NWO Vidi Grant from the Dutch Research Council (14721) to F.v.E. We also wish to thank Sisi Wang and Anna van Harmelen for their valuable input on the article.

## Data availability

All data will be made publicly available before publication.

## Code availability

Relevant code associated with the here-presented analyses will be made available through GitHub before publication.

## Supplementary Information – Figures S1-S2

**Supplementary Figure 1.**
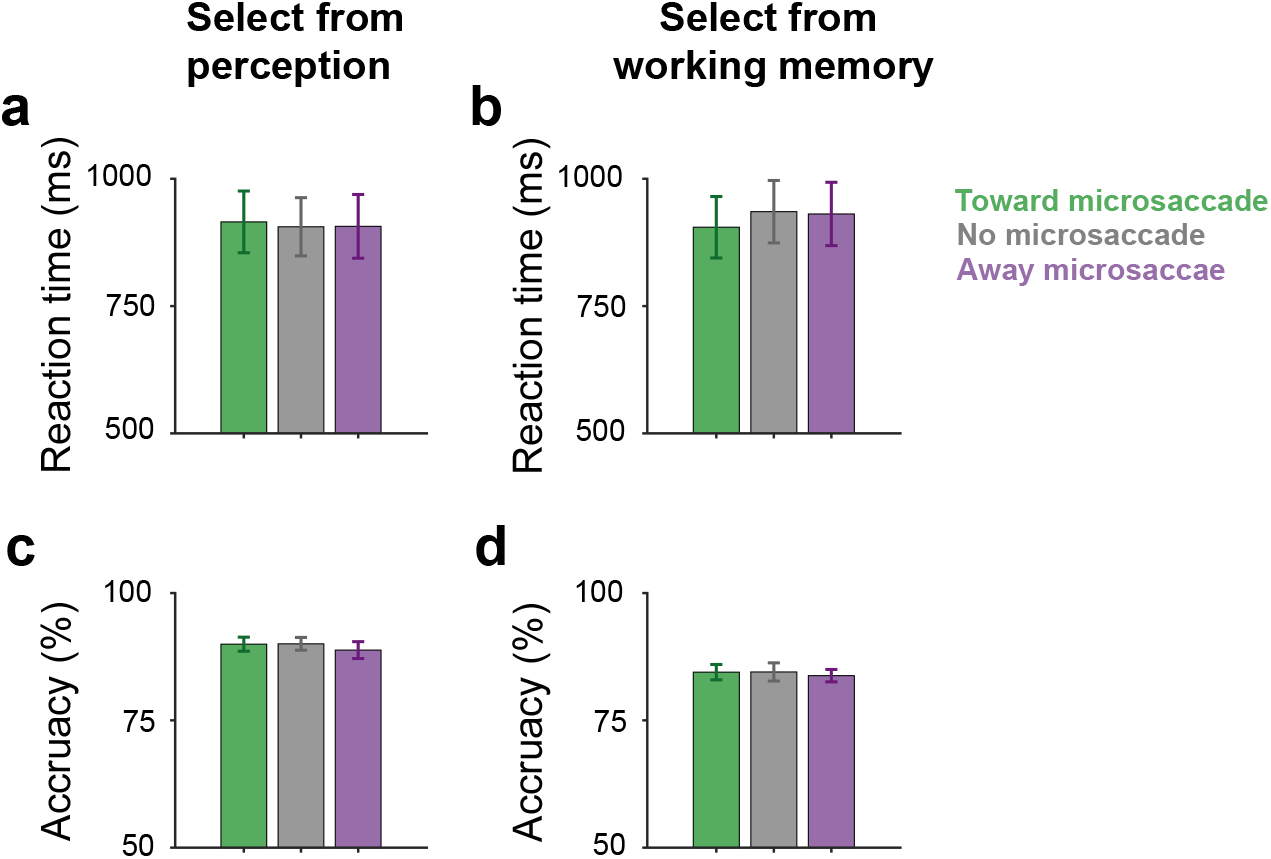
Behavioural performance as a function of microsaccade trial classes in the task requiring selection from perception (a, c) or working memory (b, d). The mean of reaction time (**a, b**) and the mean of accuracy (**c, d**). Bar graphs show mean values, with error bars indicating ±1 SEM calculated across participants (n = 23). For reaction time, a one-way ANOVA did not yield an effect of microsaccade trial-class in the perceptual-selection task (F(2, 44) = 0.2, P = 0.82, partial η2 = 0.009), but did show a main effect in the working-memory selection task (F(2, 44) = 6, P = 0.005, partial η2 = 0.21). Post-hoc t-test in the latter task showed how reaction times in toward-microsaccade trials were shorter than in both no-microsaccade (t(22) = -3.89, P_Bonferroni_ = 0.002, d = -0.81) and away-microsaccade trials (t(22) = -2.82, P_Bonferroni_ = 0.03, d = -0.59). No significant difference was found between no-microsaccade and away-microsaccade trials (t(22) = 0.4, P_Bonferroni_ = 1, d = -0.08). For accuracy, one-way ANOVAs did not yield a significant effect of microsaccade trial-class, neither in the perceptual-selection task (F(2, 44) = 1.26. P = 0.29, partial η2 = 0.05) nor in the working-memory selection task (F(2, 44) = 0.28, P = 0.76, partial η2 = 0.01).

**Supplementary Figure 2.**
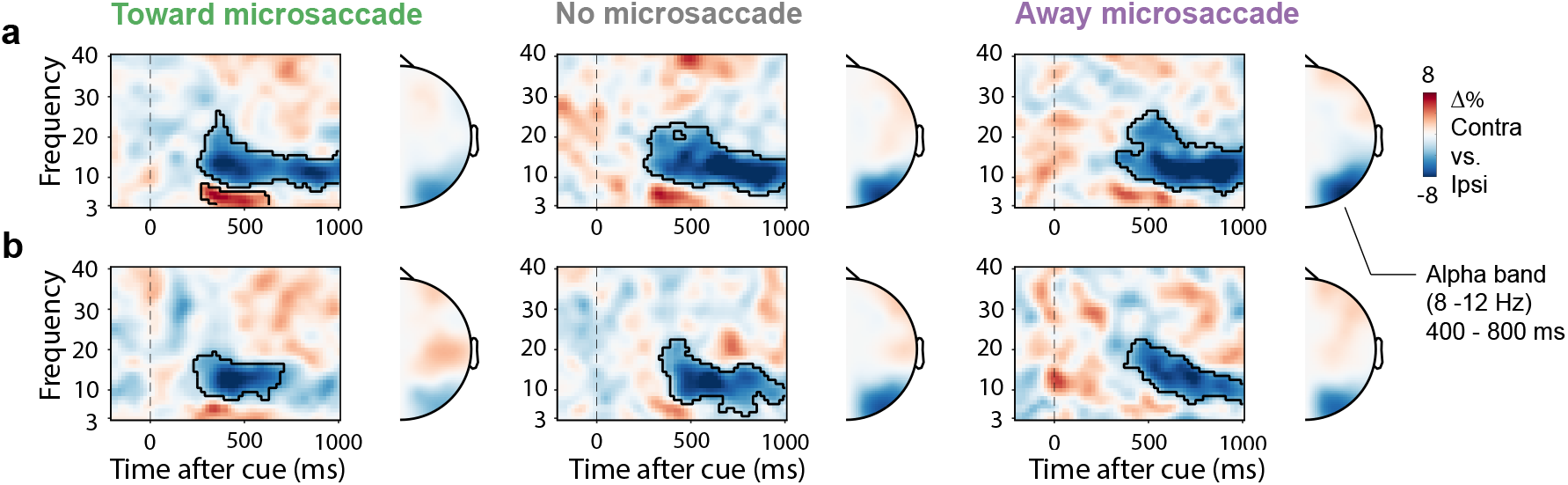
The spatial lateralization of alpha activity is clear even in the away-saccade trials despite the lack of a N2pc in these trials. **a, b**) Time-frequency maps showing lateralization of EEG activity according to the side associated with the cued visual objects when selecting from perception (panel **a**) or working memory (panel **b**). We used the same electrodes as the ones used to quantify the N2pc (PO7/8). Topographies show the lateralization of 8-12 Hz alpha-band activity in the window from 400-800 ms after cue onset (as in ^22^).

## Notes

### Competing Interest Statement

The authors have declared no competing interest.

